# Phylogeography of the pallid ground squirrel (*Spermophilus pallidicauda* Satunin, 1903) as a consequence of Holocene changes in the Mongolian steppe ecosystems

**DOI:** 10.1101/2022.11.10.515786

**Authors:** Svetlana Kapustina, Yansanjav Adiya, Elena Lyapunova, Oleg Brandler

**Affiliations:** Laboratory of Genome Evolution and Speciation, Koltzov Institute of Developmental Biology of Russian Academy of Sciences, Moscow, Russia; Institute of Biology, Mongolian Academy of Sciences, Ulaanbaatar, Mongolia

**Keywords:** speciation, steppe ecosystems, life history, Inner Asia, *Spermophilus pallidicauda*

## Abstract

The influence of Quaternary climatic changes is a source of intraspecific genetic heterogeneity of faunal components of Asian steppe and semi-desert ecosystems. The pallid ground squirrel *Spermophilus pallidicauda* is a typical representative of Inner Asian Marmotini, the intraspecific structure of which remained unstudied to date. We studied for the first time the genetic structure of the pallid ground squirrel based on cytochrome *b* and control region of mitochondrial DNA variability. We generated ecological niche models to estimate the current and past habitat suitability for *S. pallidicauda*. Our results revealed two phyletic lineages dividing this species into western and eastern population groups. According to our proposed reconstruction of the history of *S. pallidicauda* distribution, the divergence of the detected phyla may have resulted from the formation of the ecological barrier that separated the western and eastern parts of the range in the early Holocene. The hypothesis of the origin and life history of *S. pallidicauda* is given.

## INTRODUCTION

Global climate has fluctuated greatly during the past millions of years, and the inescapable consequences for most living organisms have been great changes in their distribution. It can be expected that such range changes will have genetic implications, and the advent of DNA technology provides the most appropriate markers for studying them (Hewitt 2000). Phylogeographic studies make it possible to relate the recent population structure of species to the history of their distribution due to climate change, which caused fluctuations in their ranges (Hewitt 2000; Avise 2000).

The steppe and semi-desert ecosystems of Central Asia have a long history, including rapid spatial and structural transformations under the influence of global climate changes. Past responses to climate change in the Asian steppe-desert imply that current ecosystems are unlikely to recover their present structures and biodiversity if they are transformed by human influence or severe climate change occurs because these habitats are sensitive (Barbolini *et al*. 2020).

One of the key species of Asian steppe biocenoses are ground squirrels (Thorington *et al*. 2012). In addition, they are an important object of human economic and social activity as a source of food and medicinal resources and an element of ethno-cultural heritage (Adiya 2007). Tribe Marmotini (Rodentia, Sciuridae) combines a large number of species with different structures of species ranges, as well as different levels of morphological and genetic differentiation. Specific biology features of some species of this group (dwelling in burrows, winter hibernation, sociality) caused their special sensitivity to climate changes. In this regard, ground squirrels are significantly interesting as model objects in phylogeographical research for understanding the evolution of recent biocenoses (Liapunova & Vorontsov 1970; Mantooth *et al*. 2013; Ge *et al*. 2014; Brandler *et al*. 2015a; Patterson & Norris 2016; Faerman *et al*. 2017).

Despite the high interest in Marmotini in general, the genetic variability of ground squirrels inhabiting the vast open spaces of Inner Asia (Mongolia, South Siberia, and China) remains insufficiently studied. The alternation of glacial and interglacial periods apparently led to significant changes in the location, structure, and extent of steppe ecosystems during the Quaternary (An *et al*. 2008; Khenzykhenova *et al*. 2021), which must certainly have affected widespread species with limited mobility, such as ground squirrels of the genus *Spermophilus*. The pallid ground squirrel (*Spermophilus pallidicauda* Satunin, 1903) is a typical representative of *Spermophilus* of Inner Asia and is the most eastern form of the subgenus *Colobotis* Brandt, 1844. The subgenus *Colobotis* includes the evolutionarily youngest Asian desert-steppe ground squirrels *S. major, S. erythrogenys, S. brevicauda, S. pallidicauda, S. relictus*, and *S fulvus* (Tsvirka *et al*. 2006; Kryštufek & Vohralík 2012; Kapustina *et al*. 2015a). Allopatric speciation in this group occurred due to spatial isolation formed under the influence of ecological and geographical barriers (Nikol’skii & Rumyantsev 2004). The wide area of *S. pallidicauda* extends latitudinally across the arid depressions of Mongolia from Uvs Nuur Basin on the West to Baruun-Urt Town in the East Mongolian Plain and East Gobi in Inner Mongolia, China, on the East (Bannikov 1954; Kryštufek &Vohralík 2012; Zhang 1997). It is bounded by the mountain systems of Khangai, Mongolian Altai, and Gobi Altai, Gobi Desert, and partitioned by lakes of the Great Lakes Depression and the Valley of the Lakes. The *S. pallidicauda* distribution range is geographically isolated from ranges of other *Colobotis* species (Fig. 1). Typical habitats of the pallid ground squirrel are dry fescue and wormwood steppes and feather-grass (*Stipa capillata* + *S. gobica)* steppes and gramineous-saltwort semideserts (Bannikov 1954).

**Figure 1.**
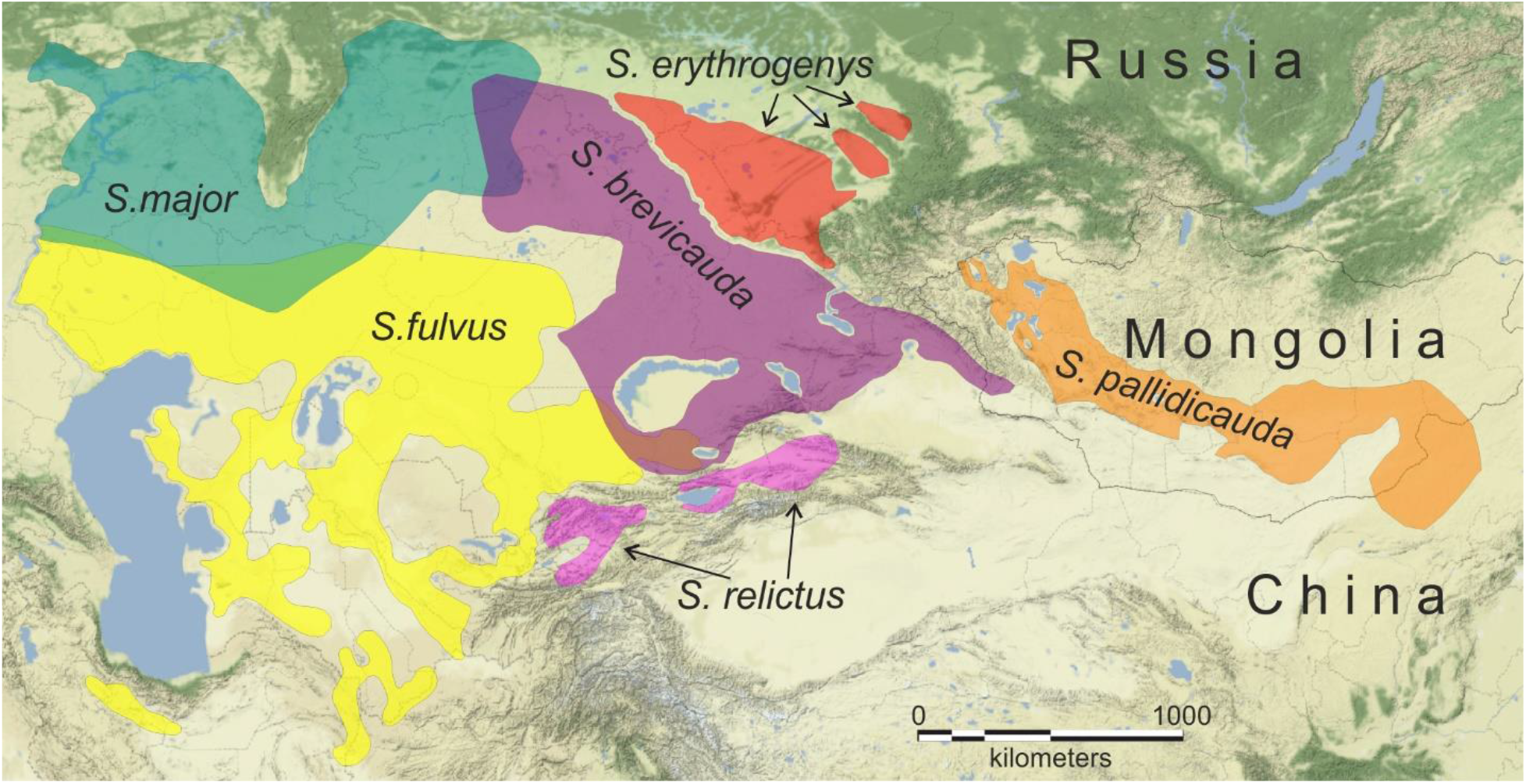
Distribution ranges of subgenus *Colobotis* species (after Ognev 1947; Bannikov 1954; Zhang et al. 1997; Kryštufek et al. 2009; Brandler et al. 2021 with our modifications).

### Taxonomic background

*Spermophilus pallidicauda* was described by K.A. Satunin (1903) from the locality of Lake Khulmu-Nur in the Mongolian Altai. Later, the species rank of *S. pallidicauda* was repeatedly questioned due to insufficiently clear morphological differentiation from other relatively close species. Ognev (1947) included this form in the Brandt’s ground squirrel (*S. brevicauda* Brandt, 1843), lowering its rank to subspecies (*S. brevicauda pallidicauda*). Later, Kuznetsov (1948) united *major*, *erythrogenys*, *pallidicauda*, and *brevicauda* forms into one polytypical species *S. major* Pallas, 1778, considering the pallid ground squirrel as a subspecies of the russet ground squirrel – *S. major pallidicauda*. Bannikov (1954), believing the *major* to be a separated species, combined red-cheeked and Brandt’s ground squirrels into the species *S. erythrogenys* Brandt, 1841, recognizing the pallid ground squirrels as its most eastern subspecies (*S. erythrogenys pallidicauda)*. This point of view remained predominant for a long time (Gromov *et al*. 1965; Kryštufek &Vohralík 2012). The species rank of S. pallidicauda was restored after describing its chromosomal set (2n=34, NF=68; Orlov *et al*. 1978), which differs from the karyotypes of other ground squirrels of the subgenus *Colobotis* (2n=36, NF=68; Vorontsov & Lyapunova 1969). Subsequently, the species independence of *S. pallidicauda* was confirmed by biochemical and molecular genetic studies (Harrison *et al*. 2003; Herron 2004; Frisman *et al*. 2014; Ermakov *et al*. 2015). The first molecular phylogeny of ground squirrels based on the variability of the cytochrome *b* mitochondrial gene (Harrison *et al*. 2003) incorporated *S. pallidicauda* with *S. alaschanicus* Büchner, 1888. This was further explained by the use of the only known hybrid *pallidicauda* X *alaschanicus* having *S. pallidicauda* mtDNA instead of *S. alaschanicus* in this study (Kapustina *et al*. 2015a, Kapustina *et al*. 2018a). Other molecular phylogenies confirmed the species independence of *S. pallidicauda* and its relatedness to *S. brevicauda* (Ermakov *et al*. 2015; Kapustina *et al*. 2015a, Matrosova *et al*. 2019).

Regardless of the taxonomic rank of the pallid ground squirrel, the intraspecific variability and population-genetic structure of this widespread form remain completely unstudied. The lack of paleontological data does not allow us to hypothesize the evolution of this species. Here we investigated for the first time the variability of two mitochondrial DNA markers from different parts of the *S. pallidicauda* distribution area to understand its intraspecific genetic structure. We conducted ecological niche modeling to assess the influence of paleoclimatic events on the formation of the range and genetic structure of *S. pallidicauda*.

## MATERIALS AND METHODS

### Tissue Sampling

Tissue samples were obtained from ground squirrels collected during our field trips in 1995, 2007, 2009-13, and 2017. The samples are deposited in the Joint Collection of Wild Animal Tissues for Basic, Applied, and Conservation Research of the Koltzov Institute of Developmental Biology RAS (IDB RAS), state registration number 6868145. In addition, three specimens from the collection of the Zoological Museum of Moscow State University (ZMMU) and one from the collection of the Zoological Institute of the Russian Academy of Sciences (ZIN) were used. We studied in total 67 individuals of *S. pallidicauda* with recorded GPS coordinates of capture points from 17 localities in Mongolia (Table 1). Twenty specimens were karyotyped earlier (Korablev *et al*. 2006) and twenty five were karyotyped for the first time using the standard technique (Ford & Hamerton 1956) to confirm species identity. To analyze intraspecific variability, local samples insignificantly distant from each other and not separated by ecological and geographic barriers were united into population samples (numbers in Table 1 and Fig. 2).

**Table 1.** List of *S. pallidicauda* specimens sequenced. Samples for which *cytb* was sequenced are marked in bold. Haplotypes of *cytb* are in parentheses. The F1 hybrid from *S. pallidicauda* female and *S. alaschanicus* male was marked in italic.

| Population | Locality | Geographical coordinates | Collection ID | CR haplotypes |
| --- | --- | --- | --- | --- |
| 1 | Uvs aimag, Olgii sum | 49.0946111°N<br>92.0276667°E | 25849, 25850, 25851, 25852, 25853, 25854, **25856(c1)**, 25857, 25858 | 1a |
|  |  |  | **25855(c1)** | 1b |
| 2 | Khovd aimag | 48.8039200°N<br>92.0522900°E | **25848(c2)** | 2a |
| 3 | Khovd aimag, Darvi sum, the right bank of the Khushatin-Gol River | 46.8833806°N<br>93.2963194°E | **25842(c3)**, 25843, 25844, 25847 | 3a |
|  |  |  | **25845(c4)**, 25846 | 3b |
|  | Khovd aimag, Darvi sum | 46.974636°N<br>93.407716°E | 25253, **25254(c5)** | 3c |
| 4 | Bayankhongor aimag, northeast of Bayan-Ondor | 44.87166667°N<br>98.95916667°E | **S-192431(c6)** | 4a |
| 5 | Bayankhongor aimag, 64 km west of Bayan-Khongor town | 46.1175°N<br>99.905833°E | ZIN 103313 | 5a |
|  | Bayankhongor aimag, 68 km west of Bayan-Khongor town | 46.133889°N<br>99.833333°E | **S-196468(c7)** | 5b |
|  | Bayankhongor aimag, Bayan-Ovoo sum | 46.0436111°N<br>100.0175°E | **25832(c8)** | 5c |
|  |  |  | **25833(c9)**, 25835, 25836, 25837, 25838, 25840, 25841 | 5d |
|  |  |  | 25834 | 5e |
| 6 | Uverkhangai aimag, 17 km south of Guchin-Us sum | 45.3123033°N<br>102.3791117°E | 25769, 25770, 25771 | 6a |
|  |  |  | 25772, **25773(c10)** | 6b |
|  |  |  | 25767 | 6c |
| 7 | Omnogovi aimag, 50 km northwest of Dalanzadgad town | 43.9°N 103.0°E | 24019 | 7a |
|  |  |  | 24018 | 7b |
| 8 | Omnogovi aimag, Gurvan-Saihan range, 15-20 km west of Dalanzadgad town | 43.5°N 104.2°E | 24022 | 8a |
|  |  |  | **24020(c11)**, 24023 | 7a |
|  |  |  | 24021 | 8b |
|  | Omnogovi aimag, Gurvan-Saihan range, 19 km west-southwest of Dalanzadgad town | 43.5282000°N<br>104.1924867°E | 25761 | 7a |
|  |  |  | 25762 | 8b |
|  | Omnogovi aimag, Gurvan-Saihan range, 15 km southwest of Dalanzadgad town | 43.4962900°N<br>104.2707100°E | 25764 | 8c |
|  |  |  | 25765, 25766 | 8d |
|  | Omnogovi aimag, northeast of Khurmen | 43.367222° N<br>104.300833°E | **S-192432(c11)** | 8e |
| 9 | Dundgovi aimag, Khuld sum, Toimert-Ula mount | 45.2603550°N<br>105.6534700°E | 25752 | 9a |
|  |  |  | 25753 | 9b |
|  |  |  | **25754(c12)** | 9c |
| 10 | Dundgovi aimag, 50 km southwest of Mandalgovi town | 45.5°N 105.8°E | 24005, 24007, 24012, 24002 | 9a |
|  |  |  | 24001, 24008, 24010, 24011, 24017 | 9c |
|  |  |  | **24004(c12)**, 24009, **24006(c12)**, 24016, | 10a |
|  |  |  | 24003 | 10b |
| 11 | Dornogovi aimag, 7 km north of the Saikhandulaan sum | 44.744148°N<br>109.047041°E | 25462 | 9a |
|  |  |  | 25463 | 11a |

**Figure 2.**
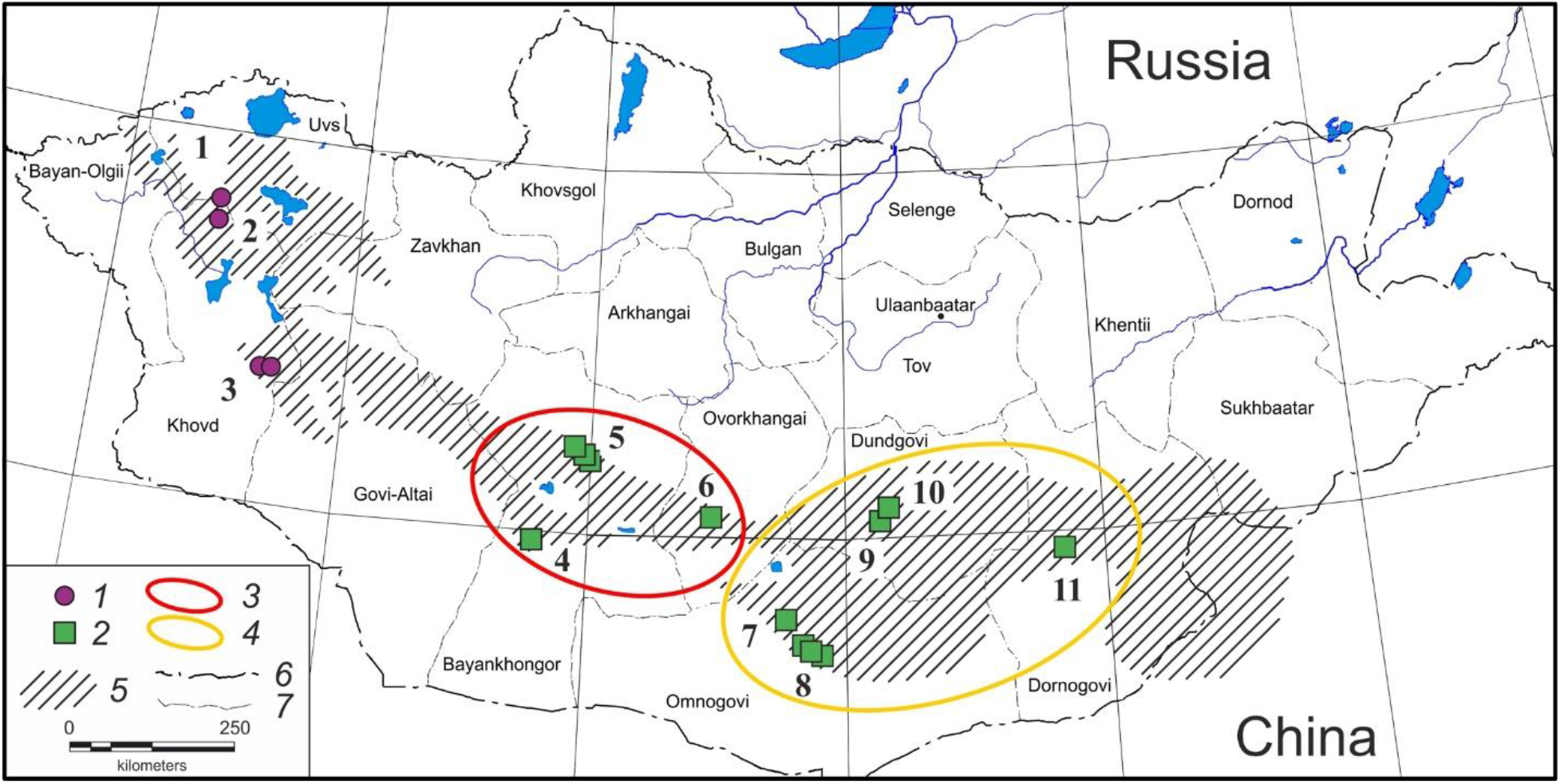
Map of localities studied for *S. pallidicauda*. Legend: 1 - populations of western phylogroup W; 2 - populations of eastern phylogroup E; 3 - population group I of phylogroup E; 4 - population group II of phylogroup E (designations of phylogroups and population groups according to cluster designations in Fig. 3, 4); 5 - distribution range of *S. pallidicauda;* 6 - Mongolian border; 7 - aimag borders. Numbers of studied localities as in Table 1.

The animals were treated according to established international protocols, as in the Guidelines for Humane Endpoints for Animals Used in Biomedical Research. All the experimental protocols were approved by the Ethics Committee for Animal Research of the Koltzov Institute of Developmental Biology RAS in accordance with the Regulations for Laboratory Practice in the Russian Federation. All efforts were made to minimize animal suffering.

### DNA Extraction, Amplification, and Sequencing

Genomic DNA was extracted by the standard salt method (Aljanabi & Martinez 1997). The nucleotide sequence of a full-length mtDNA control region (*CR*) flanked by the tRNA_Pro and tRNA_Phe genes was used as a marker of intraspecific genetic variability. In addition, the variability of the cytochrome *b* gene of mtDNA (*cytb*) was selectively studied. Modified specific primers with 5’ ends complemented by sequence primers (M13f and M13r) were used for polymerase chain reaction (PCR) (Table S1, Supporting Information). PCR was performed in 15 μl reaction mixture of Screen Mix (Evrogen, Russia) in a Veriti Thermal Cycler (Applied Biosystems, USA), at the following conditions: +95 °C – 20 sec, annealing temperature (Table S1, Supporting Information) – 40 sec, +72 °C – 40 sec (30 cycles), and final synthesis +72 °C – 10 min. The sequencing reaction was performed in 10 μl of the reaction mixture using BigDye v.3.1 kit (Applied Biosystems, USA) according to the manufacturer’s protocols and external primers: M13f and M13r (Table S1, Supporting Information). The obtained fragments were analyzed in Formamide on an ABI 3500 automatic genetic analyzer (Applied Biosystems, USA). Laboratory procedures were performed on the basis of the Core Centrum of IDB RAS.

### Molecular Data Analysis

The consensus sequences for *CR* and *cytb* were aligned with the algorithm ClustalW implemented in the MEGA X software (Kumar *et al*. 2018). For both sequence sets, HKY+G+I (Hasegawa *et al*. 1985) was selected as the best nucleotide substitution model through the Bayesian information criterion (BIC) for maximum likelihood (ML) analysis with the MEGA X. Genetic differences were assessed by pairwise distances (*p*-distance, *d_p_*). Bayesian inference analysis (BI) was performed using the default GTR+G+I model in MrBayes 3.2 (Ronquist *et al*. 2012) based on 3×10^7^ generations with each 5000th retained in two MCMC chains. Reconstruction was completed with a standard deviation of the separated frequencies of 0.002. To construct a summary tree, 90,000 of the 120,000 trees generated were used with discarding of the first 25%. The phylogenetic tree node stability was tested by bootstrap analysis with 1000 replicates (ML) and by calculating inverse probabilities in BI. Clade support was considered significant when the bootstrap value was 70% (ML) and inverse probability (BI) was 0.95.

We used sequences of *S. pallidicauda* specimens deposited in the Collection of IDB RAS (collection ID in parentheses) from GenBank NCBI: *CR* - KR611458 (25772); *cytb* - AF157866 (24004), AF157869 (24006), and AF157868 (24020) published earlier (Harrison *et al*. 2003; Brandler *et al*. 2015a) in addition to those determined by us. The *CR* sequence of *S. brevicauda* (MW149993, Brandler *et al*. 2021) and *cytb* sequences of *S. fulvus* (AF157908, Harrison *et al*. 2003) and *S. brevicauda* (MH518108, Matrosova *et al*. 2019) were used as outgroups.

The visualization of Bayesian trees was carried out in the software FigTree v1.4.3 (http://tree.bio.ed.ac.uk/). The evolutionary network of *CR* haplotypes was constructed with the Network 5.0 using the “median joining” algorithm (Bandelt *et al*. 1999).

Standard molecular diversity and neutrality test statistics was performed using the Arlequin v. 3.5.2.2 software (Excoffier & Lischer 2010) with calculation of the number of haplotypes (*H*), number of polymorphic positions (*S*), number of mutations (η), haplotype (*h*) and nucleotide (*π*) diversities, and mean number of pairwise differences (*k*). Deviations from the model of a neutrality evolution and demographical stability of population were examined using the *Fs* test (Fu 1997) basing on a coalescent simulation with 1000 repetitions.

The isolation by distance hypothesis was tested by a correlation of geographic and genetic (*F_st_*) interpopulation distances using the Mantel test in GenAlEx 6.51b2 software (Peakall & Smouse 2006; 2012) with 10,000 permutations.

### Ecological niche modeling

We generated ecological niche models (ENM) to estimate the current and past habitat suitability for *S. pallidicauda* using Maxent 3.4.4 (Phillips *et al*. 2017). Geographic records of pallid ground squirrels (records of animal or colony occurrence) collected by us (28 points) and from GBIF online databases (Global Biodiversity Information Facility; GBIF.org (23 June 2022) GBIF Occurrence Download https://doi.org/10.15468/dl.qgrchj; 23 points) were used for modeling. Geographic records were matched with the species distribution (Bannikov 1954; Sokolov & Orlov 1980; Zhang *et al*. 1997; Kryštufek & Vohralík, 2012) to detect and remove possible inaccurate records from the preliminary data sets. The final dataset had 43 occurrence records. Duplicate occurrence records were removed during model construction. Twenty percent of the presence records were reserved for tasting and eighty percent for training.

ENMs were generated for current conditions (CC; 1960-1990 u 1970-2000), the MidHolocene (MH; < 6 kya), and the Last Glacial Maximum (LGM; < 22 kya). Environmental predictors were selected from an initial set of 19 bioclimatic variables downloaded from WorldClim 2 (www.worldclim.org) and 18 bioclimatic variables downloaded from ENVIREM (ENVIronmental Rasters for Ecological Modeling, http://envirem.github.io; Title & Bemmels, 2018) related to temperature, precipitation, and topography at a spatial resolution of 2.5 arc minutes. General circulation models (GCM) CCSM4, MIROC-ESM, and MPI-ESM-P were used for MH and LGM. We restricted the calibration area for niche modeling analysis to Mongolia and the adjoining part of China with similar climatic and topographic conditions and limited by frames between 80 and 130 east longitude and 35 and 60 north latitude degrees. The contribution of each predictor was evaluated by the AUC and jackknife test, and the predictors with the highest contribution and expected greater significance for the species’ biology were selected. Next, highly correlated variables were removed using Pearson’s correlation coefficient with R-package implemented in QGIS 3.24.1 software and only variables with coefficient values smaller than 0.75 were kept. The final set of variables selected for constructing the models included 11 variables (Table S2, Supporting Information), in which only two were related to temperature; seven were related to moisture exchange, and two to topographic features.

The performance of the model selected as optimal was assessed inspecting the test of Area Under the Curve (AUC). The auto features option in MAXENT was selected to allow for linear, quadratic, product, threshold, and hinge features to describe relationships between specimen locations and environmental conditions (Merow *et al*. 2013). For model construction, we used 50,000 background points and 50,000 as the maximum number of iterations. A regularization multiplier of 2.0 was selected for calculating the mean AUC through cross validation. A total of 10 model replicates were generated for each scenario, and the median value of these 10 models was used to calculate average models. Model performance was evaluated using AUC. Raster processing and visualization were performed with MapInfo GIS software.

## RESULTS

All karyotyped ground squirrels from the total sample had chromosome sets specific to *S. pallidicauda* (2n=34) (Orlov *et al*. 1978), except for an individual previously identified as a *pallidicauda* X *alaschanicus* hybrid (2n=36) (Korablev *et al*. 2006; Kapustina *et al*. 2015a). This result confirmed the morphological identification of all the examined ground squirrels as *S. pallidicauda*.

### *CR* variability

The *CR* nucleotide sequences (1007-1011 bp) in 66 individuals of *S. pallidicauda* were determined (Table 1). The mean nucleotide composition was A = 31.1%, T = 33.6%, C = 23.3%, and G = 12.0%. The results of molecular diversity statistics and neutrality test are presented in Table 2. In the total sample, 28 unique *CR* haplotypes were recovered, in which 63 sites (6.25%) were variable, and among them 46 sites (4.56%) were parsimony-informative. The mean transition/transversion ratio was R=9.44. Each of the western populations (1-6) had a unique set of haplotypes; the southern populations 7 and 8 had one common haplotype (7a); most eastern populations (9, 10, and 11) had a common haplotype 9a (Fig. 2; Table 1).

**Table 2.** Summary of molecular diversity and neutrality test statistics by phylogroups and for the total sample of *S. pallidicauda*. Number of individuals (*N*), number of polymorphic positions (*S*), number of mutations (η), haplotype diversity (*h*), nucleotide diversity (*π*), mean number of pairwise differences (*k*), Fu’s *Fs*, standard deviation (SD).

| Samples | $N$ | $H$ | $S/\eta$ | $h \pm \text{SD}$ | $\pi \pm \text{SD}$ | $k$ | Fu's $F_s$ (p) |
| --- | --- | --- | --- | --- | --- | --- | --- |
| "Western" phylogroup | 19 | 6 | 23/23 | 0.7427±0.0852 | 0.0069±0.0038 | 6.947 | 3.96 (>0.05) |
| "Eastern" phylogroup | 49 | 22 | 33/33 | 0.9413±0.0153 | 0.0076±0.004 | 7.634 | -3.57 (>0.05) |
| Total sample | 68 | 28 | 68/70 | 0.9504±0.0109 | 0.0181±0.009 | 18.324 | 0.55 (>0.05) |

The topology of the ML and BI phylogenetic trees differed insignificantly. In both trees, haplotypes were divided into two clades (*W* and *E*, Fig. 3a) with high statistical supports. The inter-clade mean genetic distance was *d_p_*=0.0305±0.0046 (standard error (SE) hereinafter) based on 20 nucleotide substitutions. The mean within-clade distances were lower by an order of magnitude and were as follows: *W* – *d_p_*=0.0089±0.0019, *E* – *dp*=0.0074±0.0016.

**Figure 3.**
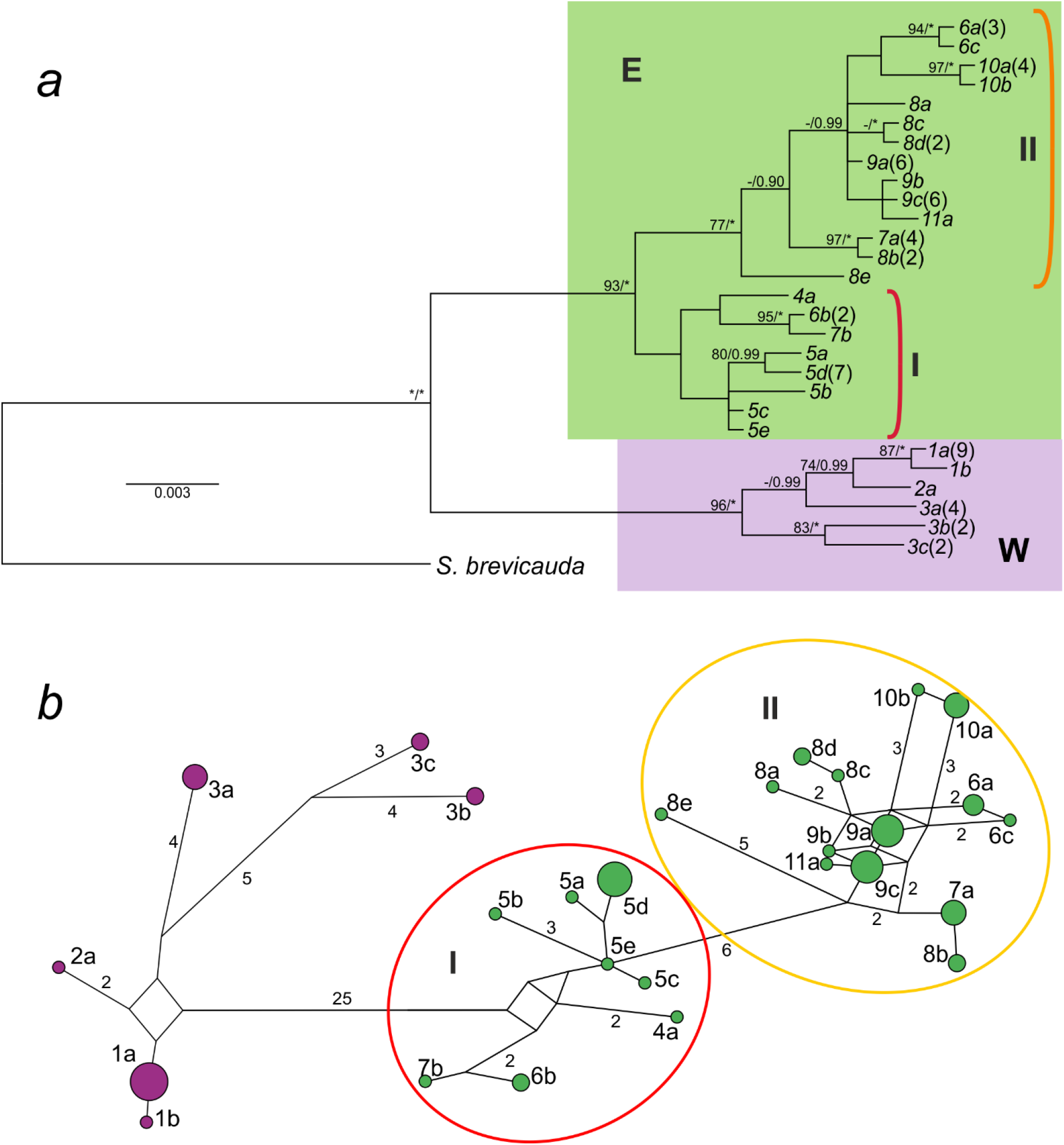
Phylogenetic tree (*a*) and median-joining network (*b*) of the observed 28 haplotypes of mitochondrial DNA control region (*CR*) sequences of *S. pallidicauda*. Numbers above branches in tree represent bootstrap support ≥70 of ML and posterior probabilities ≥0.95 of BI. Nambers near nodes in network represent numbers of substitutions. Haplotype numbers correspond to those in Table 1. Colors correspond to Fig. 2: green marks haplotypes from eastern populations (E); purple marks haplotypes from western populations (W); red and orange ovals mark haplotypes from central (I) and marginal (II) groups of eastern populations respectively.

The *W* clade included haplotypes from the western populations (1-3) (Fig. 2). It should be noted that sample 1 (Uvs aimag) was the most homogeneous one (9 out of 10 animals had a common haplotype), and sample 3 (Khovd aimag) was the most heterogeneous one (*d_p_*=0.00659±0.002, Table S3, Supporting Information). Three haplotypes from the last population were evolutionarily close but differing by 7-12 nucleotide substitutions (Fig. 3b).

The *E* clade (Fig. 3a) combined haplotypes from the populations of the central and eastern parts of the species area (4-11, Table 1), including the F1 hybrid *alachanicus* X *pallidicauda* which was caught in the Gurvan Saikhan Mountains foothill (population 8). Two lineages were integrated into this clade: one of them (*I*, Fig. 3a) contained haplotypes from localities 4-6 in the central part of the distribution (Bayankhongor, Ovorkhangai, Omnogovi aimags), while the other (*II*) united haplotypes mainly from the most eastern (7-11) localities in Omnogovi, Dundgovi, and Dornogovi aimags (Fig. 2). At the same time, haplotypes *6a* and *6c* of the eastern line *II* were found in the central population 6, and haplotype *7b* of the central line *I* was found in the eastern population 7. Haplotypes of the *E*-*II* lineage clustered irrespective of the geographic location, and formed a complex structure with many links on the evolutionary network (Fig. 3b).

Because of the detection of two phyletic lineages, we calculated molecular variability indices both for the total sample and for each lineage separately. The haplotype diversity of the total sample was high (*h*=0.9504±0.0109) (Table 2). The value *h* in group *E* was significantly higher than in *W* and was close to that of the whole sample. However, nucleotide diversities (*π*) and ranges of nucleotide variability (*k*) differed insignificantly among these groups. Low values of *π* combined with high *h* values may indicate the origin of recent populations from a few small ancestral populations over a short time. Values of the *Fs* test, although statistically insignificant, may indicate recent population growth, at least in the eastern part of the range.

The test for isolation by distance revealed a significant correlation of genetic and geographic distances within the entire range (Mantel test: *r* = 0.808, p<0.01). This correlation in the eastern part of the range was comparatively lower (*r* = 0.535, p<0.01). At the same time, this test value was insignificant, that was probably related to the small sample size, for the western part (*r* = 0.994, p>0.05), indicating that the positive correlation within the whole species was associated with the geographical disjunction of the two mitochondrial linages.

### *Cytb* gene variability

To assess the differentiation between mitochondrial lineages of *S. pallidicauda* detected by *CR* analysis, we selectively recovered the full-length *cytb* nucleotide sequences (1140 bp) for 13 individuals of *S. pallidicauda* (Table 1). In the complete sample of 16 *cytb* sequences, 12 unique haplotypes were detected, in which 20 sites were variable (1.75%), and among them 14 sites were parsimony informative (1.23%). The average nucleotide content was A = 28.6%, T = 34.1%, C = 24.8%, G = 12.5%. The mean genetic distance within the total sample was *d_p_*=0.0077±0.002. The haplotype diversity (*h*) of the general sample was 0.9583±0.0363, and nucleotide diversity *π*=0.0053±0.003. Values of the Fu’s test *Fs* were not significant.

The phylogenetic tree constructed based on *cytb* gene variability (Fig. 4) did not contradict the tree based on *CR* variability. The sample of *S. pallidicauda* was split up into two well-supported clusters. Cluster *W* included haplotypes from the western part of the area, and cluster *E* combined haplotypes of specimens from the eastern part. In the latter cluster, central (4-6, haplotypes *c6-c10*) and eastern (8-10, *c11* and *c12*) populations were divided into two clusters (Fig. 4). The composition of the identified clusters fully corresponded to that found with our *CR* tree analysis. The *cytb* haplotype divergence between *W* and *E* clusters was *d_p_*=0.0074±0.0020 (0.74%). The average genetic distance within cluster *W* (*d_p_*=0.0027±0.0012) was lower than in *E* (*d_p_*=0.0052±0.0017) in contrast to nearly equal average genetic distances within these clusters founded by us with *CR*.

**Figure 4.**
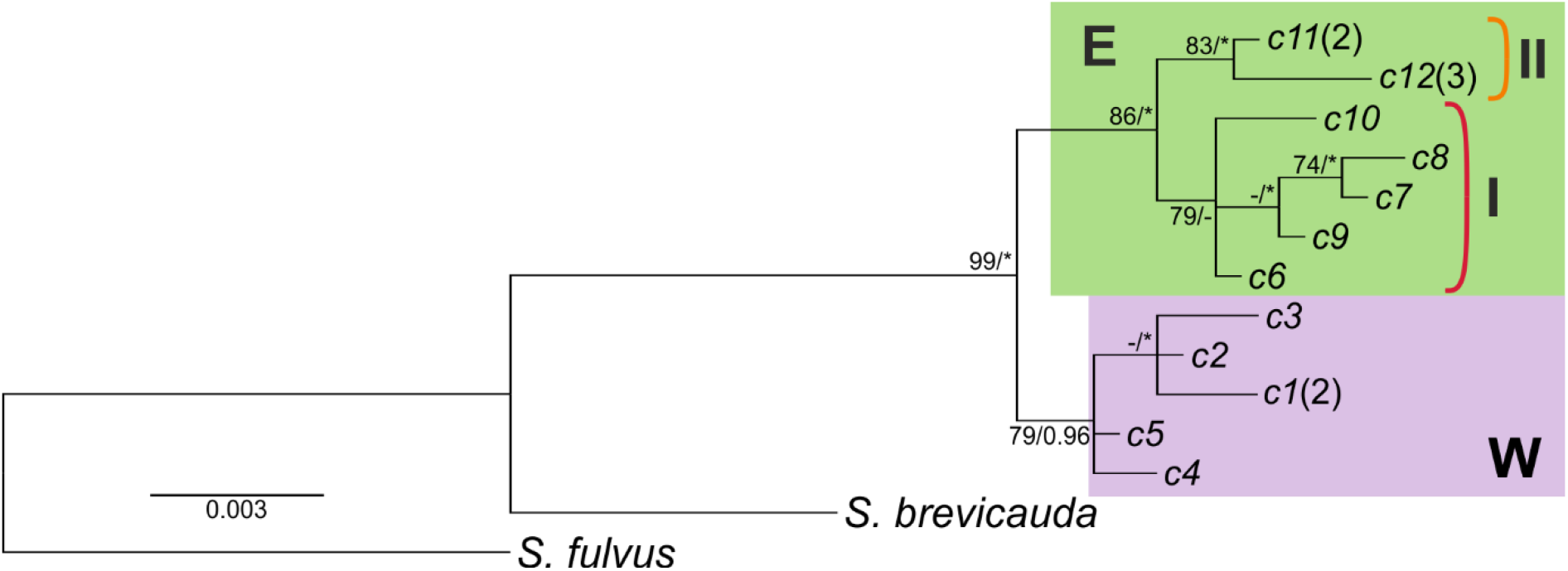
Phylogenetic tree of the observed 12 haplotypes of mitochondrial cytochrome *b* gene (*cytb*) sequences of *S. pallidicauda*. Numbers above branches in tree represent bootstrap support ≥70 of ML and posterior probabilities ≥0.95 of BI. Haplotype numbers correspond to those in Table 1. Colors correspond to Fig. 3.

All new DNA haplotype sequences were deposited in GenBank under accession numbers OP530507-OP530543.

### Models of *S. pallidicauda* distribution

The average test AUC for the 10 replicates was 0.982, and the standard deviation was 0.011, indicating that model prediction accuracy was greater than random. The recent distribution of suitable abiotic conditions closely corresponded to the known present-day distribution of *S. pallidicauda* (Fig. 5a). This result indirectly confirms the species status of the pallid ground squirrel, exposing a combination of factors which characterizes the ecological niche of this species as different from other closely related ones. The model showed no significant gap in suitable abiotic conditions over the range between the identified *E* and *W* groups of populations.

**Figure 5.**
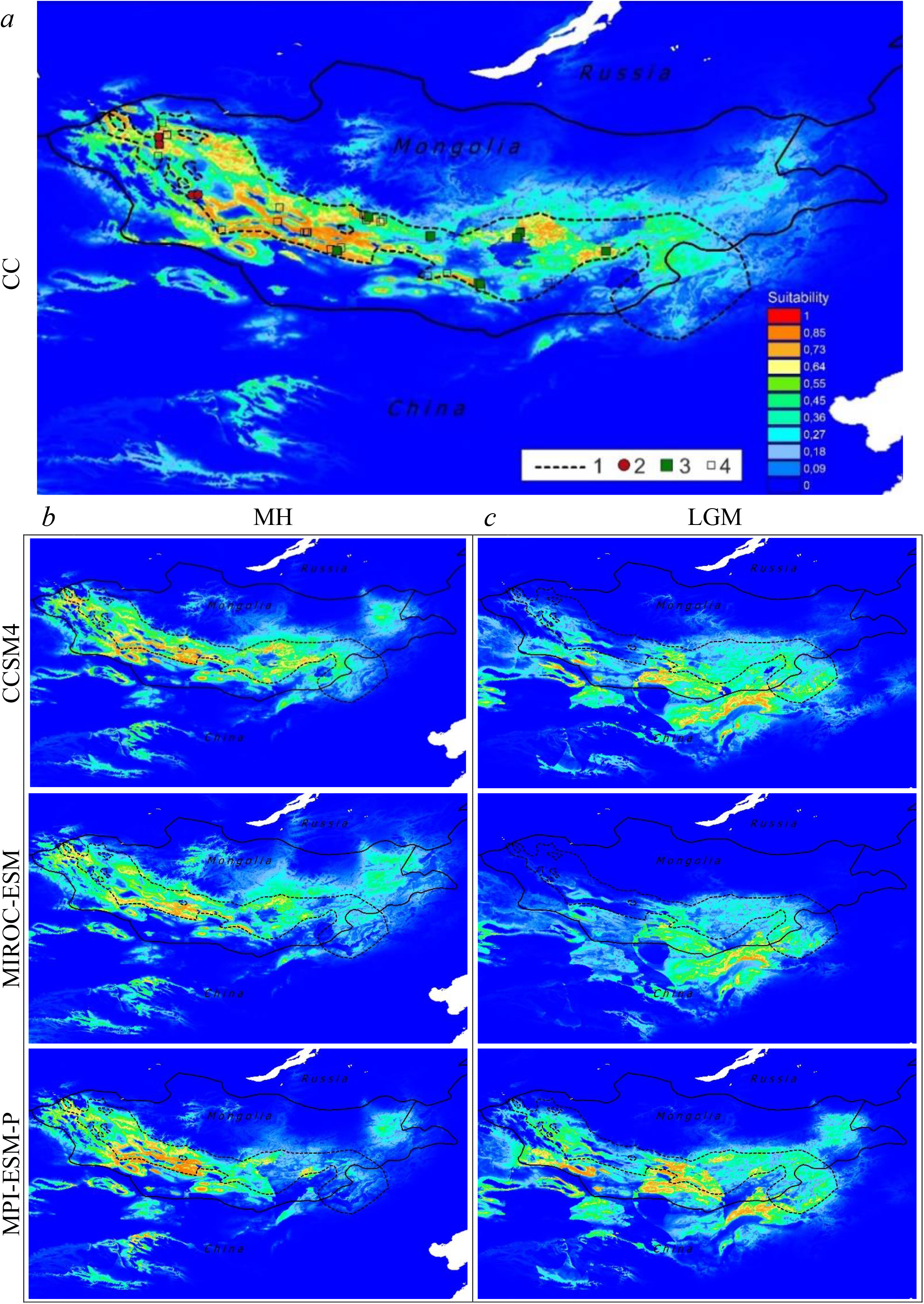
Ecological niche models of *S. pallidicauda* for the present (*a*), Mid-Holocene (*b*), and Last Glacial Maximum (*c*). Warmer colors indicate higher suitability for species occurrence, as depicted in the key. Legend: 1 – *S. pallidicauda* current distribution; 2 – populations of western phylogroup W; 3 - populations of eastern phylogroup E; 4 – other points of occurrence records of *S. pallidicauda*. CCSM4, MIROC-ESM, and MPI-ESM-P are general circulation models used.

Spatial projections from the estimated climatic scenarios showed a shift of areas with suitable habitat conditions from the south and southeast in the north and west directions from the late Pleistocene to the Mid-Holocene. The possible distribution during the LGM may have consisted of one or two parts, depending on the GCM applied. The main part was located eastward in Inner Mongolia and eastern Gobi, and the second part comprised some isolated refugia in the southern foothills of the Mongolian Altai (Fig. 5c). Unsuitable conditions for the species were present in the southern Mongolian Altai, Valley of the Lakes, and Goby Altai areas according to all climatic scenarios. By the MH, the spatial distribution of conditions suitable for the species approximated to the current state. (Fig. 5b). The most comfortable climatic conditions were formed in the center of the recent *S. pallidicauda* range in the Valley of the Lakes and intermountain depressions between the southern Mongolian Altai and Khangai, while the eastern part of the range was the least suitable. At the same time, a suitable habitat for the species had almost completely disappeared in Inner Mongolia.

## DISCUSSION

In our study, we present the first range-wide assessment of genetic structure for the pallid ground squirrel, predominantly inhabiting Mongolia. Based on mitochondrial DNA data, our analyses revealed that *S. pallidicauda* populations are divided into two main lineages. The intraspecific variability of *S. pallidicauda* shows a clear geographic localization of haplotypes and genetic diversity of individual populations.

### Taxonomic implications

The *S. pallidicauda* was considered by all authors to be monophyletic. However, we have for the first time revealed its intraspecific heterogeneity associated with geographical variability. The average *cytb* sequence divergence between the *W* and *E* phylogroups (0.77%) is similar to the difference between European ground squirrel’s subspecies *S. citellus citellus* and *S. c. karamani* (0.98%) (Kryštufek *et al*. 2009; Kryštufek &Vohralík 2012). It was previously noted that mean sequence divergences are low between species in the subgenus *Colobotis* (Kapustina *et al*. 2015a). For example, the divergence by *cytb* of 2.9%-3.9% between the phyletic lineages “brevicauda”, “intermedius”, and “iliensis” of *S. erythrogenys* s. l. allowed the authors to suggest their raising to species rank (Matrosova *et al*. 2019).

Comparison of *CR* variability of the pallid ground squirrel and other species of the genus *Spermophilus* is also ambiguous. The *CR* sequence divergence between cryptic species (former subspecies) of speckled ground squirrels *S. suslicus* (2n=34) and *S. odessanus* (2n=36) is much higher (4.4%) (Brandler *et al*. 2015a). The differences between *S. pallidicauda* phylogroups are twice as great as the differentiation of the phyletic lineages (presumably subspecies) of *S. dauricus* (*d_p_*=0.015) (Kapustina *et. al*. 2018b) and similar to the differences between “eastern” and “western” phylogroups of *S. major* (0.8%) (Brandler *et. al*. 2021).

The lack of an analysis of intraspecific morphological variation of the pallid ground squirrel and poor development of a common subspecies criterion for genetic differentiation leave open the question of the taxonomic significance of the *S. pallidicauda* phyletic lineages we have detected. However, the above comparisons with other studies of intraspecific variability in *Spermophilus* may indicate the potential possibility of subspecies division of this taxon.

### Life history of *S. pallidicauda*

The low level of intraspecific genetic variability, high *h* values, and low *π* values found for *S. pallidicauda* as a whole indicate a relatively short evolutionary history of this species accompanied by a significant reduction and fragmentation of the range in the recent past. The results on the *CR* molecular variability allow us to suggest that the origins of the western and eastern populations representing the two phyletic lineages have followed similar scenarios but have some specific details.

The populations of the western part of the range seem to have undergone a significant decline in abundance and passed through a “bottleneck” recently. This is indicated by the relatively low value of *h* and small *π* (Table 2), low haplotype diversity, and the absence of common haplotypes for populations of the western phylogroup (*W*). This conclusion is also supported by the high value of the Mantel test, which indicates the current isolation and lack of gene flow between the Uvs aimag (1 and 2, Fig. 2) and Khovd aimag (3) populations. The current spatial isolation of western populations was evidenced by our 2011 surveys, as a result of which we found no ground squirrels or their burrows in the area between population 3 and Khar-Us-Nuur Lake. Our observations were consistent with the result of ENM of the current range (Fig. 5a).

For populations from the central (Bayankhongor and Ovorkhangai aimags) and eastern (Omnogovi, Dundgovi, and Dornogovi aimags) parts of the range, a high *h* level in combination with a low *π* level, as well as the *Fs* test result indicate population expansion from a few isolated localities in the recent past. The division of the *E* phylogroup into two branches indicates the existence in the past of a disjunction of central and most eastern populations of *S. pallidicauda*. However, the presence of both central and eastern haplogroups in populations 6 and 7 indicates a recent migration exchange between the eastern and central partitions of the range.

The question of the history of the pallid ground squirrel’s range formation is very poorly developed. Fossils that could be classified to this species or its ancestral form are extremely rare. There are only two known *S. pallidicauda* fossils, but detailed information on them is not available in the research literature (Gromov *et al*. 1965). Given the relatively low intraspecific level of genetic differentiation of this species, we assume that the events that caused this kind of differentiation took place in the relatively recent past. Here we propose a hypothesis of historical changes in the range of *S. pallidicauda* in the Late Pleistocene and Holocene based on genetic data, ENM results, and literature paleoclimatic data.

*Spermophilus pallidicauda* is most closely related both phylogenetically and geographically to the Central Asian species *S. brevicauda*, the genetic distance from which in *cytb* is 2.7% (Kapustina *et al*. 2015a). These two species differ well in karyotype (2n=34 vs. 2n=36) and their ranges are allopatric (Kryštufek & Vohralík 2012). Such a ratio of chromosomal and molecular differences is typical for chromosomal speciation, some scenarios of which involve the fixation of chromosomal rearrangements in isolated populations. (Vorontsov 1960; White 1978; Bakloushinskaya 2016). At present, the ranges of these species are separated by the Mongolian Altai Mountains. However, it is admissible to assume the existence of a common ancestral area in the past. The area of the modern habitat of *S. pallidicauda* had a wetter climate until the end of the Pleistocene, clearly not suitable for arid species. At that time, in the Great Lakes Depression and the Valley of the Lakes, paleolakes (with an area of more than 90,000 km^2^ in the first) were formed in the Middle and early Upper Pleistocene including the fifth marine isotope stage (MIS 5) (Yang *et al*. 2004). It can be assumed that the *brevicauda*-*pallidicauda* ancestral range could have formed after that in the late Pleistocene in areas located south of the modern *S. pallidicauda* range, including the southwestern macroslopes of the Mongolian Altai and Goby Altai and northern parts of the Gobi Desert, where steppe biocenoses were distributed during MIS 3 (Lehmkuhl *et al*. 2018; Yu *et al*. 2019). The ensuing cooling and the development of extremely arid climatic conditions throughout Central Asia led to the development of glaciers and cold deserts (Lehmkuhl & Haselein 2000; Yang *et al*. 2004), while the tundra dominated in the intermountain depressions and desert landscapes of Mongolia (Khenzykhenova *et al*. 2021). The ENM reconstructions for LIG indicated the absence or dramatic reduction of areas suitable for *S. pallidicauda* habitat in the western near Altai regions of Mongolia and China at this time (Fig. 5c). At the same time, suitable ecological conditions persisted in the south of Mongolia and in Inner Mongolia. Possibly, the formation of the 34-chromosome form *S. pallidicauda* adapted to dwelling in the extremely continental arid conditions of Central Asia took place at this time in isolation from the main range of red-cheeked ground squirrels. This conclusion is consistent with the low level of genetic divergence between *S. pallidicauda* and *S. brevicauda*, which indicates a relatively short period of their independent evolution.

Throughout the Holocene, cyclical changes of wetter and drier climatic periods may have caused repeated changes in the number, structure, and boundaries of the range of the pallid ground squirrel. The invasion of *S. pallidicauda* from the east into the intermountain depressions of northwestern Mongolia could probably have occurred during the Early Holocene, when most of the lakes became shallow to the maximum and dry and desert steppes prevailed there. (Khenzykhenova *et al*. 2021). At the same time, the expansion of the Gobi Desert should have shifted the range of this species northward to the Gobi Altai. At the end of this period, an increase in humidity began, which continued for the first half of the Mid-Holocene until about 6 kyr BP (An *et al*. 2008). Lake levels had risen at this time, the forest and forest-steppe were spread out much further south than they are today, and deserts were transformed into steppe landscapes (Khenzykhenova *et al*. 2021). Taiga forests were widespread in the Khangai, Mongolian, and Goby Altai until 4.5-3.5 kyr BP (Tarasov *et al*. 2000; Khenzykhenova *et al*. 2021). Given this, it is possible to assume a significant reduction in desert-steppe landscapes suitable for *S. pallidicauda* habitation in the space between the Great Lakes Depression and the hilly plains of Uverkhangai during the maximum humid phase (about 8-7 kyr BP) due to the shortening distance between forested mountains and moistening of intermontane depressions between 95° and 100° E. We can date the divergence of the western (*W*) and eastern (*E*) phyletic lineages of *S. pallidicauda* and the fragmentation of the western part of its range to isolated populations at this time. Subsequent cooling and increase in aridity combined with an increased climate continentality led to a gradual restoration of steppe and semi-desert biocenoses in the intermountain Altai-Khangai depressions by 4000 yrs BP (An *et al*. 2008; Khenzykhenova *et al*, 2021). The ENM reconstructions (Fig. 5b) showed the development of the most suitable habitat conditions for *S. pallidicauda* in the central part of the modern range, as well as a continued isolation and fragmentation of western populations and a significant fragmentation or complete disappearance of eastern populations in the Mid-Holocene. This may be due to the unevenness of climatic changes, as well as an earlier aridization of eastern and southern parts of Mongolia compared to the northwestern part (An *et al*. 2008). The division of the phylogroup *E* into two subgroups (Fig. 3, 4) may indicate the maintenance of small relict ground squirrel populations in the east during this period, the small size of which is evidenced by low π values. These refugia (or one of them) may have become a source of westward expansion as the area habitable for the species expanded. Such an expansion of eastern populations, confirmed by the Fs-test values, could have occurred in the Late Holocene (<3000-2000 yrs BP) when the eastern part of Mongolia was covered by xerophytic steppes (Khenzykhenova *et al*. 2021) and might have been supported by a slight increase in humidity around 1500 yrs BP (An *et al*. 2008). The finding of central and eastern subgroup haplotypes in the populations of Gurvan Saikhan and southeastern foothills of Khangai (populations 6 and 7) indicates the association region of ranges of the central and eastern subgroups during the species areal recovery. Apparently, such gene flow could have occurred periodically during the formation of suitable conditions in the intermountain Khangai-Gobi Altai depressions and eastward on the undulating-hilly plains. At present, such links are probably limited due to the lack of suitable ecological conditions (Fig. 5a).

Recent historical changes in the range of *S. pallidicauda* are apparently associated mainly with local changes in habitat conditions. In conditions of sufficient stability of steppe biocenoses and their insignificant disturbance by human activity, global climate changes in the region have the greatest influence on the dynamics of the pallid ground squirrel habitat, both in the past and at present. For example, pallid ground squirrels were abundant in the Dornogovi, Dundgovi, and Omnovi aimags in 1995, but their numbers in this region declined dramatically by 2010-2014 under the influence of unfavorable climatic changes, particularly long-term droughts (Brandler *et al*. 2015b, our unpubl. data).

The differentiation of *S. pallidicauda* into eastern and western phylogroups appears to be caused by past fragmentation of the range under the influence of paleoclimatic events. A similar differentiation within the range extended in the latitudinal direction was found in a number of species of the steppe and open space mammalian fauna of Mongolia. In ground squirrels, division into two phyletic lineages based on genetic markers was found in the Mongolian marmot *Marmota sibirica*, Radde, 1862 (Kapustina *et al*. 2015b) and long-tailed ground squirrel *Urocitellus undulatus*, Pallas, 1778 (Vorontsov *et al*. 1978; Kapustina *et al*., 2014; McLean *et al*., 2018). At the same time, the level of genetic differences in both species is significantly greater than in the pallid ground squirrel, which indicates more ancient paleoclimatic processes fragmenting steppe biocenoses similarly. Among other steppe species, latitudinal genetic subdivision was found in *Microtus mongolicus*, Radde, 1861 (Bannikova *et al*. 2019), *Lasiopodomys brandtii*, Radde, 1861 (Li *et al*. 2017), and *Procapra gutturosa*, Pallas, 1777 (Sorokin & Kholodova 2006). The level of molecular-genetic differentiation of the eastern and western groups of populations in these species is somewhat different, but unequivocally indicates the existence of a paleogeographic barrier or a few barriers, possibly repeatedly formed in the Central Mongolian regions in the Middle or Upper Pleistocene. The different values of the described genetic differences can be associated both with uneven evolutionary rates and different durations of isolation, as well as with specifics habitat preferences and different levels of migration potential (for example, *P. gutturosa* has much higher migration activity compared to territorially conservative rodents). Determination of the influence of paleoclimatic events on the genetic diversity of steppe mammals of Inner Asia requires further studies of their genomes and comparative analysis with paleoclimatic and paleogeographic data.

## Conclusion

Our results revealed the genetic heterogeneity of *S. pallidicauda*, which is considered a monotypic species. Evaluation of the taxonomic significance of its two spatially separated phylogroups requires further research at the morphological level. A solution to this problem can also contribute to the development of genetic criteria for the subspecies category.

The evolution of the pair of species *S. brevicauda* - *S. pallidicauda* appears to be a good model of allopatric chromosomal speciation. It can be assumed that the formation of *S. pallidicauda* as a separate species took place due to a fragmentation of the peripheral parts of the ancestral range under the influence of powerful paleoclimatic changes, which could have led to an isolation and further divergence of the 34-chromosomal form. The spatial and temporal parameters of range changes in such a process of speciation are not sufficiently clear at the present time. Our attempt to reconstruct the history of development of the ancestral range based on ecological niche modeling agreed well with the observed genetic diversity. However, a description of the earlier events that accompanied the separation of the 34 chromosome *S. pallidicauda* from the 36 chromosome *S. brevicauda* may be possible only if we analyze genetic data as well as range reconstructions for both species.

A comparative analysis of the genetic structures of open space species in Mongolia and adjacent territories indicates periodic fragmentations of steppe territories by ecological barriers. This is reflected in the existence of “western” and “eastern” phyletic lineages in a number of species. The dating of genetic divergences and reconstruction of paleoareas of such species could significantly contribute to our understanding of the evolution of open space biocenoses of Inner Asia.

## Acknowledgments

The authors sincerely thank the leadership and staff of the Joint Russian-Mongolian Complex Biological Expedition of the Russian Academy of Sciences and Mongolian Academy of Sciences for their help in organizing field research. We thank V.S. Lebedev, who kindly shared samples of *S. pallidicauda* from the ZMMU and ZIN collections to be included in this study. We thank D.M. Schepetov for laboratory advice and for assistance with the molecular work. The research was conducted using equipment from the Core Centrum of the Koltzov Institute of Developmental Biology RAS.

