## Supplementary material for "Phylogeography of the pallid ground squirrel (*Spermophilus pallidicauda* Satunin, 1903) as a consequence of Holocene changes in the Mongolian steppe ecosystems": Table S1

***Supporting Information***

Table S1. List of used primers for amplification and sequencing reactions.

| Purpose | Primer name | Nucleotide sequences 5’-3’ | Annealing temperature | Source |
|---|---|---|---|---|
| PCR *CR* | MDL | TCCACCTTCAACTCCCAAAGC | 62°C | Ermakov *et al.* 2002 |
| | H00651 | TAACTGCAGAAGGCTAGGACCAAACCT | | Kocher *et al.* 1989 |
| PCR *cytb* | L14725 | TGAAAAAYCATCGTTGT | 48°C | Steppan *et al.* 1999 |
| | H15915 | TCTTCATTTYWGGTTTACAAGAC | | Harrison *et al.* 2003 |
| Sequencing reaction | M13f | TGTAAAACGACGGCCAGT | 48°C | Ekimova *et al.* 2015 |
| | M13r | CACAGGAAACAGCTATGACC | | Ekimova *et al.* 2015 |
