## Supplementary material for "Phylogeography of the pallid ground squirrel (*Spermophilus pallidicauda* Satunin, 1903) as a consequence of Holocene changes in the Mongolian steppe ecosystems": Table S2

***Supporting Information***

Table S2. The final set of variables selected for ecological niche modeling

| Variable abbreviation | Brief description | Database |
|---|---|---|
| BIO1 | Annual Mean Temperature | WorldClim 2 |
| BIO12 | Annual Precipitation | WorldClim 2 |
| BIO15 | Precipitation Seasonality (Coefficient of Variation) | WorldClim 2 |
| BIO17 | Precipitation of Driest Quarter | WorldClim 2 |
| aridityIndexThornthwaite | Thornthwaite aridity index: Index of the degree of water deficit below water need | ENVIREM |
| climaticMoistureIndex | A metric of relative wetness and aridity | ENVIREM |
| growingDegDays5 | Sum of mean monthly temperature for months with mean temperature greater than 5℃ multiplied by number of days | ENVIREM |
| PETDriestQuarter | Mean monthly potential evapotranspiration (PET) of driest quarter | ENVIREM |
| PETseasonality | Monthly variability in potential evapotranspiration | ENVIREM |
| topoWet | SAGA-GIS topographic wetness index | ENVIREM |
| tri | Terrain roughness index | ENVIREM |
