## Supplementary material for "Phylogeography of the pallid ground squirrel (*Spermophilus pallidicauda* Satunin, 1903) as a consequence of Holocene changes in the Mongolian steppe ecosystems": Table S3

***Supporting Information***

Table S3. Mean genetic intra- and inter-population distances ($d_p$) of *S. pallidicauda* based on mtDNA control region (*CR*) variability data, SE – standard error

| Population | N | Intra-population ($d_p$±SE) | Inter-population ($d_p$ – below the diagonal, SE – above the diagonal) | | | | | | | | | | |
|---|---|---|---|---|---|---|---|---|---|---|---|---|---|
| | | | 1 | 2 | 3 | 4 | 5 | 6 | 7 | 8 | 9 | 10 | 11 |
| 1 | 10 | 0.0000±0.000 | | 0,0022 | 0,0023 | 0,0051 | 0,0052 | 0,0049 | 0,0049 | 0,0051 | 0,0051 | 0,0051 | 0,0052 |
| 2 | 1 | - | 0,0030 | | 0,0031 | 0,0054 | 0,0055 | 0,0053 | 0,0053 | 0,0055 | 0,0055 | 0,0055 | 0,0056 |
| 3 | 8 | 0.0066±0.002 | 0,0063 | 0,0093 | | 0,0052 | 0,0053 | 0,0050 | 0,0050 | 0,0052 | 0,0052 | 0,0052 | 0,0053 |
| 4 | 1 | - | 0,0179 | 0,0208 | 0,0212 | | 0,0020 | 0,0026 | 0,0027 | 0,0032 | 0,0030 | 0,0031 | 0,0031 |
| 5 | 11 | 0.0017±0.001 | 0,0185 | 0,0215 | 0,0219 | 0,0034 | | 0,0025 | 0,0026 | 0,0031 | 0,0030 | 0,0031 | 0,0031 |
| 6 | 6 | 0.0024±0.001 | 0,0174 | 0,0203 | 0,0207 | 0,0055 | 0,0061 | | 0,0017 | 0,0013 | 0,0009 | 0,0011 | 0,0012 |
| 7 | 2 | 0.0074±0.003 | 0,0186 | 0,0216 | 0,0220 | 0,0067 | 0,0074 | 0,0037 | | 0,0017 | 0,0017 | 0,0017 | 0,0018 |
| 8 | 10 | 0.0024±0.001 | 0,0186 | 0,0216 | 0,0217 | 0,0076 | 0,0083 | 0,0031 | 0,0046 | | 0,0009 | 0,0011 | 0,0012 |
| 9 | 3 | 0.0000±0.000 | 0,0179 | 0,0208 | 0,0212 | 0,0060 | 0,0066 | 0,0015 | 0,0037 | 0,0016 | | 0,0007 | 0,0008 |
| 10 | 14 | 0.0015±0.001 | 0,0189 | 0,0219 | 0,0223 | 0,0070 | 0,0077 | 0,0026 | 0,0043 | 0,0027 | 0,0011 | | 0,0011 |
| 11 | 2 | 0.0015±0.002 | 0,0186 | 0,0216 | 0,0220 | 0,0067 | 0,0074 | 0,0022 | 0,0045 | 0,0024 | 0,0007 | 0,0018 | |
